# Long-term memory performance optimization via Neural network-based curve fitting in *Drosophila*

**DOI:** 10.64898/2026.06.22.733713

**Authors:** Yung-Ching Lu, Chih-Ying Chen, Ling-Hui Yen, Chi-Lien Yang, Ya-Ding Liu, Wen-Jun Chen, Kuan-Lin Feng, Ming-Chin Wu, Ann-Shyn Chiang, Da-Jeng Yao, Chi-Ming Ho, Shih-Hwa Chiou, Li-An Chu

**Affiliations:** Department of Biomedical Engineering and Environmental Sciences, National Tsing Hua University; Hsinchu 300044, Taiwan; Department of Medical Research, Taipei Veterans General Hospital; Taipei 11217, Taiwan; Institute of Biotechnology, National Tsing Hua University; Hsinchu 300044, Taiwan; Institute of NanoEngineering and MicroSystems, National Tsing Hua University; Hsinchu 300044, Taiwan; Brain Research Center, National Tsing Hua University; Hsinchu 300044, Taiwan; Institute of Biophotonics, National Yang Ming Chiao Tung University; Taipei 11221, Taiwan; Institute of Systems Neuroscience and Department of Life Science, National Tsing Hua University; Hsinchu 300044 Taiwan; Kavli Institute for Brain and Mind, University of California San Diego; La Jolla, CA 92093-0526, USA; Institute of Physics, Academia Sinica; Taipei 11529, Taiwan; Department of Biomedical Science and Environmental Biology, Kaohsiung Medical University; Kaohsiung 80780, Taiwan; Institute of Molecular and Genomic Medicine, National Health Research Institutes; Miaoli 35053, Taiwan; Graduate Institute of Clinical Medical Science, China Medical University; Taichung 404327, Taiwan; Department of Power Mechanical Engineering, National Tsing Hua University; Hsinchu 300044, Taiwan; Mechanical and Aerospace Engineering Department, University of California, Los Angeles, Los Angeles, California, 90049, U.S.A.; Institute of Pharmacology, National Yang Ming Chiao Tung University; Taipei 112304, Taiwan; Genomics Research Center, Academia Sinica; Taipei 115201, Taiwan; National Yang Ming Chiao Tung University Hospital; Yilan 260006, Taiwan; Department of Biomedical Engineering, National Taiwan University; Taipei 106216, Taiwan

**Author notes:** Corresponding author. Email: Li-An Chu,; Shih-Hwa Chiou.

## Abstract

Long-term memory (LTM) formation typically requires extensive training or highly salient experiences, limiting learning efficiency. Operant conditioning is generally thought to produce stronger memory than classical conditioning because of its active learning component. Unexpectedly, however, laser-based social conditioning in *Drosophila melanogaster* revealed that while classical paradigms yielded lower short-term memory (STM) scores but higher LTM retention, operant paradigms exhibited higher STM scores followed by rapid LTM decay. To resolve this discrepancy, we employed the AI Complex Systems Response (AI-CSR) framework, which reconstructs high-dimensional learning landscapes from sparse sampling and predicts globally optimal training conditions. Following AI-CSR optimization, operant conditioning produced a twofold increase in LTM scores, yielding the strongest 24-hour social memory performance reported in flies to date and revealing the expected superiority of active learning, which had conversely shown poorer performance under standard training protocols. In contrast, AI-CSR did not further enhance classical conditioning performance but reduced training time by 50%. Single-cell RNA sequencing revealed expanded neuronal recruitment marked by the activation and inhibition of various gene combinations. Together, these findings link circuit-level reorganization with molecular programs underlying efficient long-term memory and demonstrate how AI-guided optimization can uncover latent learning capacity in biological systems.

## Main

Long-term memory (LTM) formation in humans and other animals typically requires repeated training or highly salient experiences ^1-7^. In *Drosophila melanogaster*, a widely used model for memory studies, LTM is usually induced through multiple training cycles ^3,8-10^. For example, olfactory classical conditioning requires 5–10 spaced training cycles, each pairing odor exposure with electric shock, to generate memories lasting beyond 24 hours ^8^; inter-trial intervals are essential for effective LTM formation. In contrast, operant conditioning in *Drosophila* has been studied primarily in courtship suppression ^11,12^, where repeated rejection by mated females over prolonged training periods (typically 6–10 hours) leads to persistent suppression of courtship behavior.

Building on our previously developed ALTOMS system ^13^, which decomposes social memory into classical and operant components, we examined how experimental parameters shape learning outcomes. We uncovered a striking dissociation between classical and operant conditioning. Classical conditioning scores peaked across a broad parameter range, whereas operant conditioning—particularly long-term memory (LTM)—was confined to a narrow parameter window. Within this window, flies initially exhibited lower learning scores but showed markedly enhanced performance after 24 hours. Operant LTM scores were the highest among all paradigms tested, reaching levels comparable to those typically observed in short-term memory (STM).

Efficient learning has long been a central goal in education and neuroscience. Learning protocols depend on multiple interacting variables—including duration, intensity, repetition, and spacing— creating a vast search space. Despite their widespread use, protocol design still lacks a systematic optimization framework; conventional trial-and-error approaches are time-consuming and typically converge to local optima. Drawing on complex systems science, we discovered the AI Complex Systems Response (AI-CSR) equation, a general governing equation that resolves system responses across the full parameter domain. Combined with established links between molecular and cellular mechanisms—such as synaptic connectivity, protein synthesis, and RNA expression ^14-16^—AI-CSR translates mechanistic insights into actionable strategies for memory enhancement ^17^ .

## Results

### Laser-Based Conditioning Reveals Convergent Long-Term Memory States

We established classical and operant conditioning paradigms compatible with the ALTOMS system to train male *Drosophila melanogaster* (Fig. 1A). In classical conditioning, the conditioned stimulus was the presence of a female, and the unconditioned stimulus was repetitive laser punishment delivered at fixed intervals and intensity. This design parallels olfactory classical conditioning, in which electric shocks are repeatedly paired with odors ^8^.

**Fig. 1.**
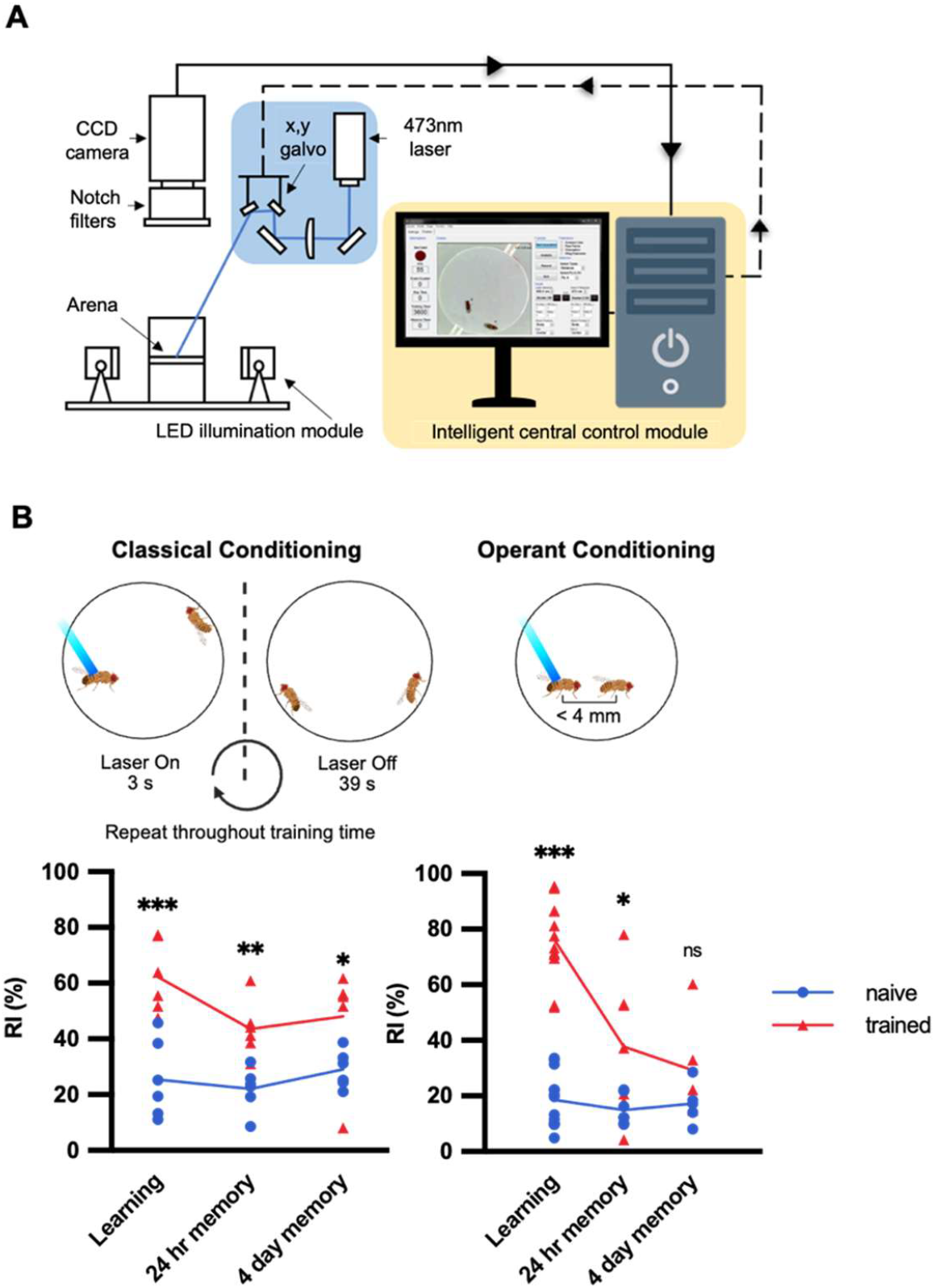
Create classical and operant social memory with ALTOMS. (**A**) ALTOMS setup. (**B**) Definitions and behavioral outcomes of classical and operant conditioning. Males were trained for 3600 s under three experimental conditions. In classical conditioning, males experienced the same laser schedule in the presence of a female. In operant conditioning, males received 20 mW laser punishment only when approaching females within 4.5 mm. Behavioral performance was evaluated immediately after training, and at 24 hr, and 4 day post-training. Data are presented as mean ± SEM. Statistical significance was determined using unpaired two-tailed Student’s t-test (n = 5–15 flies per group). ns, not significant; *p < 0.05; **p < 0.01; ***p < 0.001.

Classical conditioning elicited a robust immediate memory response, with the restraining index (RI) reaching ∼60, but declined and stabilized at ∼40 from 24 hours to 4 days post-training (Fig. 1B). In contrast, operant conditioning used the same conditioned stimulus (female fly) but delivered laser punishment contingent on male behavior, specifically inter-fly distance. To avoid punishment, males had to cease approaching and actively retreat. This action–outcome contingency produced rapid learning, consistent with prior work ^13^, yielding acquisition-phase RI values near 80 within 1 hour. Unexpectedly, RI declined to ∼40 after 24 hours—matching classical conditioning—and remained stable for up to 4 days (Fig. 1B).

This pattern has traditionally been attributed to a ceiling in *Drosophila* long-term memory, with RI values plateauing near 40, discouraging further optimization. We instead hypothesized that this limit reflects suboptimal training conditions rather than neural capacity. For laser-based conditioning, outcomes depend on multiple parameters—including laser power, punishment bout length, and training duration—and their respective levels, which have been selected largely by heuristic trial and error. With *P* parameters each tested at *L* levels, the resulting *L*^P^combinatorial space renders exhaustive testing impractical.

### Operant Conditioning Achieves Maximal Long-Term Memory Enabled by AI-CSR Equation

Biological systems often involve high-dimensional parameter spaces in which interacting variables give rise to complex systems. These systems consist of many interacting components that self-organize into dynamic, nonlinear behavior with emergent properties. Conventional reductive reasoning cannot directly infer emergent system behavior from individual components because of intricate self-organization processes.

Using an inductive reasoning approach, we discovered the Complex Systems Response (CSR) equation, which directly links emergent system properties to interacting components ^17^. The CSR equation is a second-order nonlinear polynomial, implying a single global optimum in the emergent response (Equation 5). The unknowns are the (*P*^2^ + 3*P* + 2)/2 response coefficients; thus, only (*P*^2^ + 3*P* + 2)/2 ≪ *L*^P^tests are required to reconstruct the CSR surface across the full parameter domain^17,18^.

To determine whether *Drosophila* long-term memory can be further enhanced through parameter optimization, we investigated three parameters for operant conditioning (training duration, restraining distance, and laser power) and four parameters for classical conditioning (training duration, bout length, total punishment duration, and laser power). While these models theoretically require 10 and 15 coefficients respectively, we conducted 13 and 17 experimental tests to ensure robust estimation. These parameter ranges were defined through preliminary experiments (Extended Data Fig. 1) to ensure efficacy while avoiding disruption of natural fly behavior. The resulting AI-CSR response surfaces and predicted global optima are shown in Fig. 2A.

**Fig. 2.**
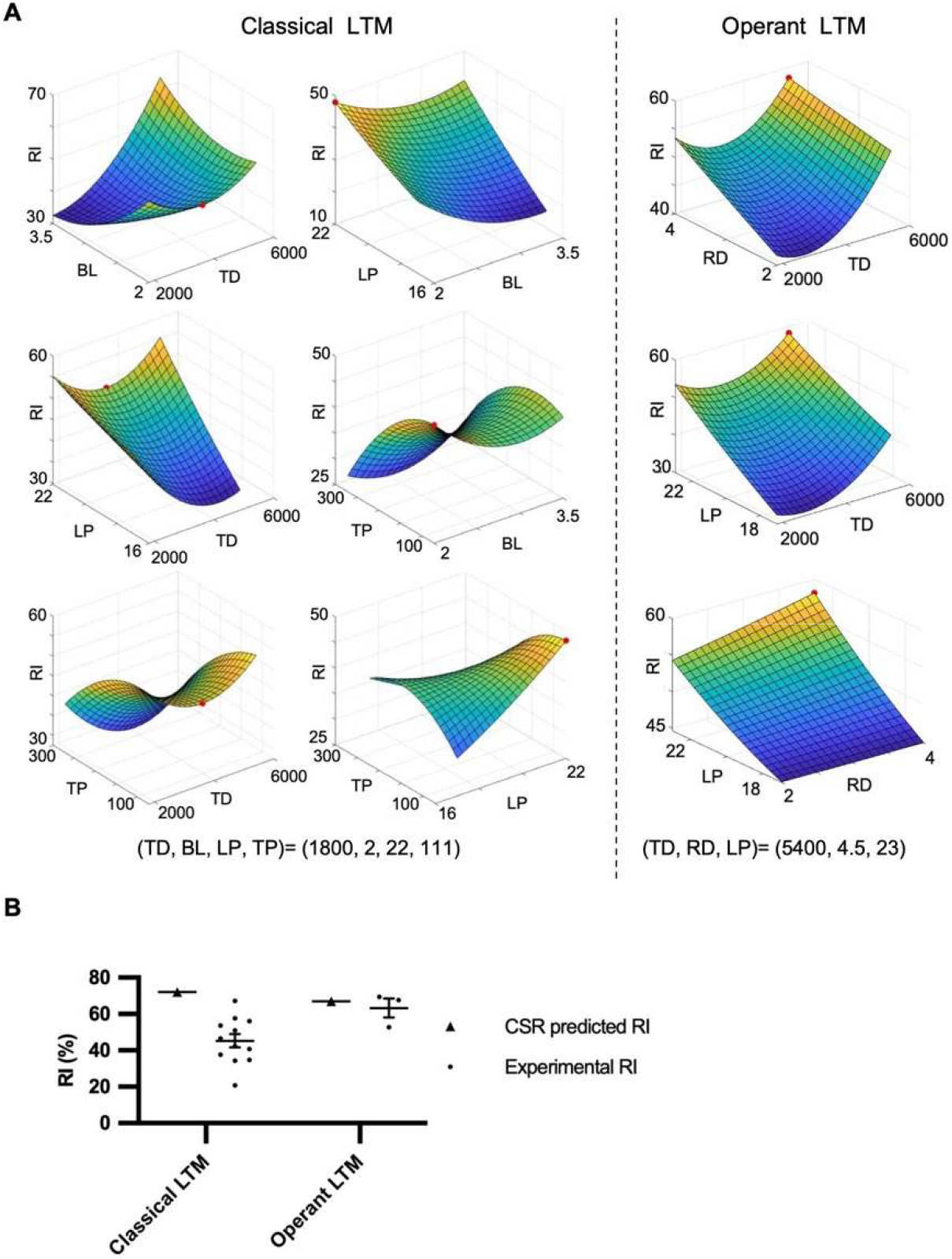
Optimization of classical and operant social memory training using AI-CSR. (**A**) The complex system response (AI-CSR) framework was applied to identify optimal parameter combinations for social memory training. Each response surface is a visualization of the relationship between the two process parameters in the parameter space. Experimental results were fitted using polynomial functions to generate a 3D response surface, with the red dot indicating the predicted global optimum. The specific values of the optimized parameter combinations derived from the CSR fitting are detailed below the panels. (**B**) Comparison of predicted and experimental outcomes across the LTM conditions, demonstrating that CSR reliably forecasts behavioral performance. Data are presented as mean ± SEM.

To identify optimal conditions for long-term memory (24 h), data from both paradigms were analyzed using the AI-CSR equation to reconstruct multidimensional response surfaces. (Supplementary Equation 1 and 2). For classical conditioning, four parameters—training duration (TD), laser power (LP), bout length (BL), and total punishment duration (TP)—were modeled using 17 parameter combinations (Supplementary Table 1). AI-CSR surface revealed a single global maximum (Fig. 2A) with a predicted restraining index (RI) of 72.0, which was experimentally validated with an RI of 45.2 (Fig. 2B). For operant conditioning, three parameters—TD, LP, and restraining distance (RD)—were analyzed across 13 combinations (Supplementary Table 2). The AI-CSR equation predicted an RI of 66.9 (Fig. 2A), well above the baseline (∼40), and was validated by a mean experimental RI of 63.3, the highest 24-hour memory score reported in *Drosophila* assays (Fig. 2B).

Beyond optimizing long-term memory (LTM) for classical and operant conditioning individually, we examined the temporal effects of AI-CSR–optimized parameter sets. The optima for classical and operant conditioning (CLS-LTM Opt and OP-LTM Opt, respectively) were applied outside their intended time points to assess memory retention. In classical conditioning, parameter variation had little effect on memory: all conditions produced robust RI values (60–80 immediately after training and 40–50 at 24 h). Thus, CLS-LTM Opt primarily improves training efficiency—for example, reducing training duration from 3,600 to 1,800 s— without altering ultimate memory outcomes (Fig. 3A). In contrast, operant conditioning showed a striking temporal dissociation. Flies trained with the original protocol and tested at 24 h exhibited RI values near 40, and flies trained with OP-LTM Opt showed similarly low performance immediately after training. Remarkably, OP-LTM Opt produced maximal performance at 24 h, revealing a delayed enhancement effect (Fig. 3B). This finding challenges the assumption that optimal LTM training also maximizes short-term memory and suggests that the mechanisms underlying LTM formation differ fundamentally from those supporting short-term memory.

**Fig. 3.**
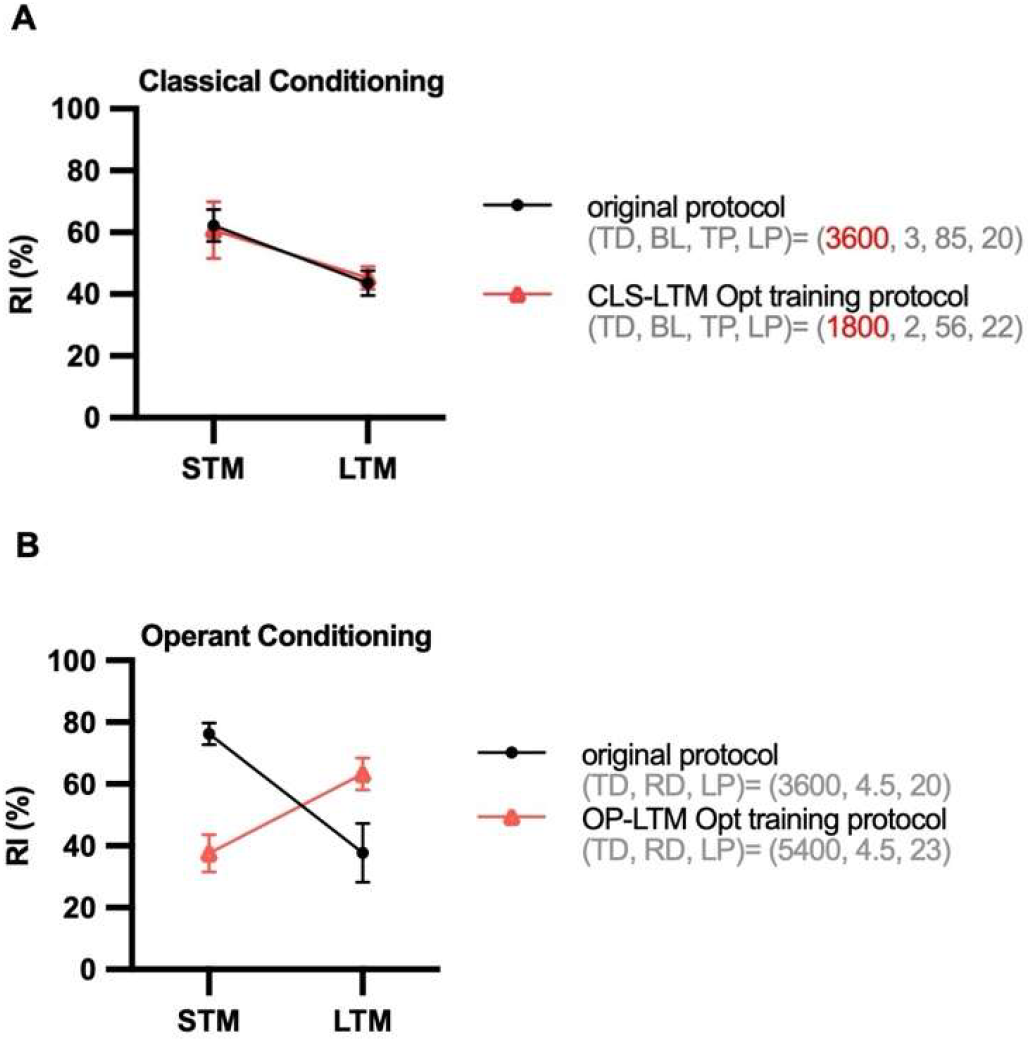
Comparison of memory performance across training protocols. Two training protocols were evaluated for each paradigm: the original regime (●) and the CSR– optimized LTM regime (△). Parameter sets for each protocol are listed in parentheses: classical (TD, BL, LP, TP); operant (TD, RD, LP). Memory was assessed immediately after training (STM) and 24 hours later (LTM), with performance quantified as the Restraining Index (RI; mean ± SEM). (**A**) Classical conditioning. Both protocols yielded comparable memory performance. The original protocol (STM: n = 6; LTM: n = 6) and the CLS-LTM Opt protocol (STM: n = 6; LTM: n = 12) showed similar scores across both short-term and long-term intervals. (**B**) Operant conditioning. Distinct memory dynamics were observed between the two training regimes. The original protocol (STM: n = 17; LTM: n = 7) followed a typical memory decay pattern. In contrast, the OP-LTM Opt protocol induced only modest STM (n = 3) but exhibited enhanced memory retention at 24 hours (n = 3).

### Single-Cell Analysis of *Hr38* Activation Reveals Neuronal Networks Underlying Optimized Long-Term Memory

To probe the molecular and cellular mechanisms underlying optimized social memory, we applied AI-CSR–derived LTM parameter sets to perform single-cell RNA sequencing (scRNA-seq) of adult *Drosophila* brains. After quality control filtering to remove background, ambient RNA, and doublets (Supplementary Figure 2), we analyzed 61,923 cells (Control, 13,347; CLS-LTM Opt, 24,630; OP-LTM Opt, 23,946), which were assigned to 216 clusters based on differential gene expression. Comparison with published markers ^19-21^ identified 70 neuronal cell types (Fig. 4, A and B). Cholinergic (VAChT), glutamatergic (VGlut), and GABAergic (Gad1) neuron proportions were comparable across groups, indicating consistent sampling (Fig. 4C).

**Fig. 4.**
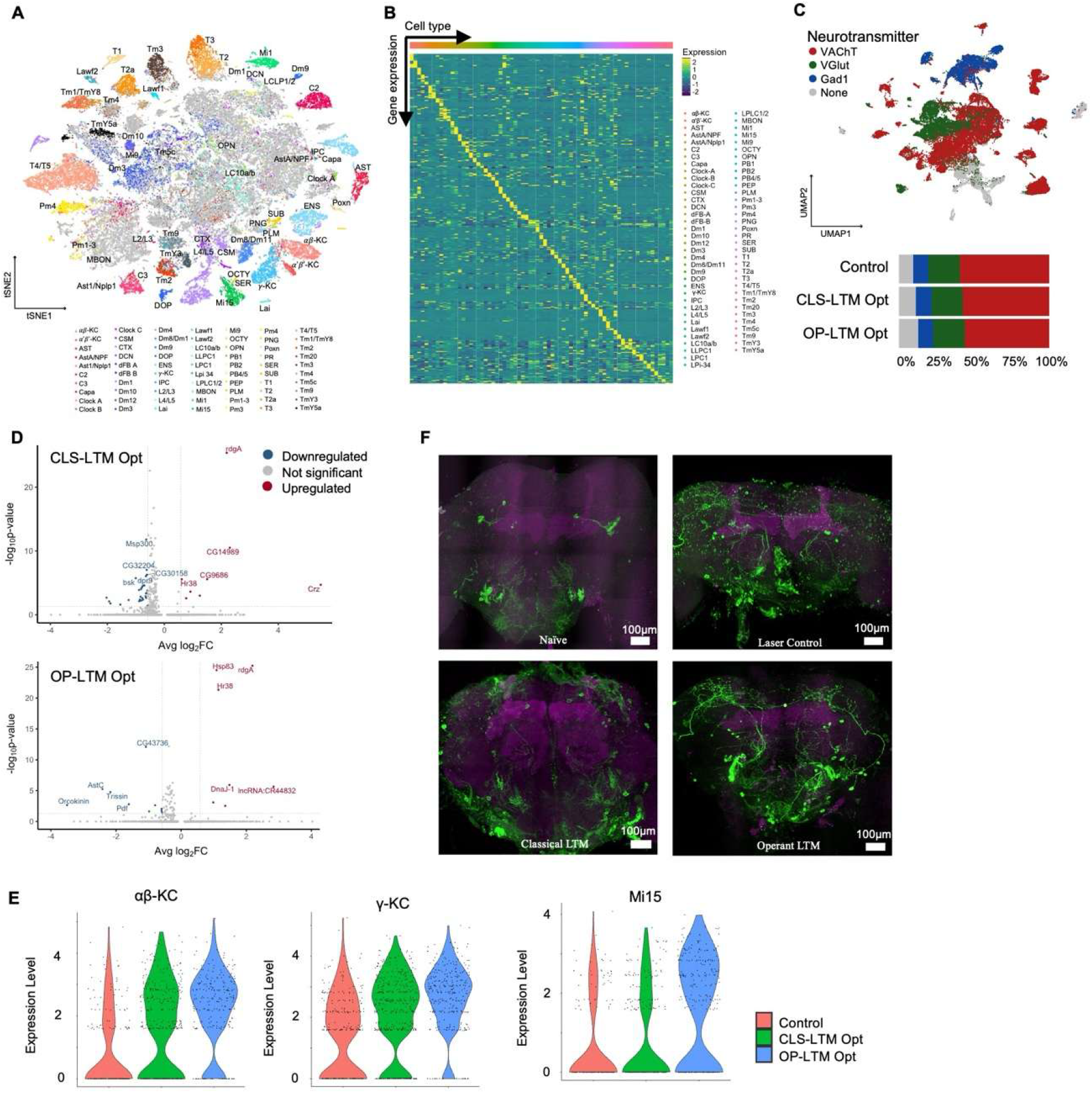
The single-cell transcriptomics analysis of adult Drosophila brains with various training conditions. (**A**) The visualization of cell type classification and annotation in adult *Drosophila* brains. (**B**) The distinct gene expression of each cell type in adult Drosophila brains. (**C**) The proportion of neurotransmitter-expressing neurons, including cholinergic (*VAChT*; red), glutamatergic (*VGlut*; Green), and GABAergic (Gad1; blue) neurons, in control, CLS-LTM Opt, and OP-LTM Opt flies. (**D**)The differentially expressed gene (DEG) analysis was used to identify downregulated (blue) and upregulated (red) DEGs in each training group compared to the control group. (**E**)Violin plots illustrate the log-normalized expression levels of *Hr38* in αβ -KC, γ-KC, and Mi15 cells across three groups: Control (red), CLS-LTM Opt (green), and OP-LTM Opt (blue). Each dot represents an individual cell, and the width of each violin reflects the kernel density estimation of the cell population at a given expression level. (**F**)Anatomical validation of Hr38-positive neurons (Hr38-Gal4>UAS-GFP, green; neuropil marker DLG, magenta). Maximum intensity projections of brains from naïve, laser control (males were placed alone in the arena and exposed to a 20 mW laser with a 3 s on/39 s off cycle), CLS-LTM Opt, and OP-LTM Opt–trained flies expressing UAS-GFP under Hr38-Gal4 control show enhanced GFP signals following OP-LTM Opt training, especially in the mushroom body and superior dorsofrontal protocerebrum (SDFP).

Differential gene expression analysis revealed training-dependent transcriptional changes (p < 0.05, |fold change| > 1.5; Fig. 4D and Extended Data Fig. 3). Consistent with recent findings identifying the immediate early gene *Hr38* as an essential transcription factor for LTM consolidation in mushroom body neurons ^22^, we observed that Hr38 was strongly induced in γ and αβ Kenyon cells (KCs) at 24 h after both CLS-LTM Opt and OP-LTM Opt training. Notably, OP-LTM Opt additionally elevated *Hr38* expression in six neuronal populations, including Mi15, LC10a/b, DOP, and TM9 neurons (Extended Data Fig. 3). Direct comparison showed higher *Hr38* levels in αβ-KC, γ-KC, and Mi15 under operant versus classical conditions (Fig. 4E), indicating broader neuronal recruitment during optimized operant memory formation. To validate the anatomical distribution of *Hr38*-expressing neurons, we used Hr38-Gal4 to drive UAS-GFP expression and compared CLS-LTM Opt– and OP-LTM Opt–trained brains with laser-only controls. OP-LTM Opt training produced distinct GFP-positive clusters in the superior dorsofrontal protocerebrum and surrounding the mushroom body (Fig. 4F). This expanded activation pattern suggests that highly efficient operant training recruits a broader spatial network of neurons, consistent with enhanced long-term memory performance.

To examine cell–cell interactions underlying differences between long-term (LTM) and short-term memory (STM) formation, we predicted ligand–receptor interactions—focusing on the Wnt signaling pathway—among αβ-, α′β′-, and γ-Kenyon cells (KCs), mushroom body output neurons (MBONs), AstA/NPF neurons, and dopaminergic neurons (DOPs) under CLS-LTM Opt and OP-LTM Opt conditions using FlyPhoneDB ^23^. Wnt signaling is known to play a key role in LTM formation ^24^. Notably, Wnt-related ligands were selectively expressed by MBONs in OP-LTM Opt flies and were predicted to interact with multiple neuronal populations, including KCs, AstA/NPF, and DOP neurons (Extended Data Fig. 4a). This Wnt-mediated communication was absent in CLS-LTM Opt, indicating a specific role for MBON-driven Wnt signaling in optimized LTM formation. Gene expression analysis further showed that the Wnt ligand *Wnt5* was specifically upregulated in MBONs of OP-LTM Opt flies, whereas Wnt receptors *Arrow* (*Arr*) and *Frizzled* (*Fz* and *Fz2*), together with the downstream effector β-catenin (*Arm*), were elevated in AstA/NPF neurons only under OP-LTM Opt conditions (Extended Data Fig. 4b). These results suggest activation of canonical Wnt signaling from MBONs to AstA/NPF neurons during operant LTM consolidation.

## Discussion

Despite these advances, several limitations remain. First, although *Hr38* is a robust marker of neuronal activation, it does not establish functional necessity for memory storage. Future studies using genetic silencing or activation of *Hr38*-expressing neurons will be required to test causal links between circuit engagement and behavior. Second, although we mapped the spatial distribution of activated neurons, the underlying mechanisms likely involve complex spatiotemporal dynamics. Memory consolidation may depend on sequential recruitment of multiple neuronal ensembles rather than sustained activity within a fixed population. Thus, artificial activation of these neurons alone may be insufficient to recapitulate the memory engram; nonetheless, our study provides a foundational map of gene regulatory networks associated with enhanced behavioral performance.

Recent single-cell transcriptomic studies have highlighted the transcriptional control of LTM, identifying the immediate early gene Hr38 as an essential transcription factor within mushroom body (MB) neurons during traditional courtship conditioning ^22^. Consistent with their findings, our scRNA-seq results confirm the pivotal role of Hr38 activation in MB γ and αβ Kenyon cells during LTM formation. However, while previous observations describe an MB-specific transcriptional trace, our AI-CSR-optimized operant paradigm (OP-LTM Opt) reveals a profound spatial expansion of this memory network. We demonstrated that under highly efficient operant training, Hr38 activation extends well beyond the canonical MB circuits to encompass a distributed neuronal population, including Mi15, LC10a/b, DOP, and TM9 neurons.

Furthermore, distinct from traditional conditioning paradigms, our systems-level approach uniquely uncovered the operant-specific engagement of MBON-driven Wnt signaling to AstA/NPF neurons. Together, these findings challenge the assumption of a static memory capacity in Drosophila and suggest that optimal training parameters not only enhance behavioral performance but achieve this by recruiting a broader, specialized brain-wide molecular and cellular network.

## Materials and Methods

### *Drosophila* strains and rearing conditions

All *Drosophila melanogaster* strains were maintained under 40–50% relative humidity on a 12:12 h light:dark cycle. Flies were cultured on standard cornmeal–agar medium (per liter: 100 g anhydrous *D*-glucose, 47.27 g organic maize flour, 25 g autolyzed yeast, 7.18 g agar, and 12.18 g Tegosept dissolved in 8.36 ml absolute ethanol). All flies used for behavioral conditioning were 5–7 days post-eclosion and group-housed (∼20 individuals per vial) after eclosion.

For behavioral conditioning assays, male flies carrying *UAS-mCD8::GFP* (BDSC #5137) and *VT64246-Gal4* (VDRC #264246), as well as wild-type *w[1118]* (BDSC #3605) females, were reared at 25 °C.

For imaging experiments, males carrying *Hr38-Gal4, tub-Gal80*^*ts*^; *UAS-mCD8::GFP* (*w[*]; P{w[+mC]=IT*.*GAL4}Hr38[02306-G4], P{w[+mC]=tubP-GAL80[ts]}[[10 / CyO; P{w[+mC]=UAS-mCD8::GFP*.*L}LL6*) were obtained from the Kyoto *Drosophila* Stock Center (DGRC #118610). Wild-type *w*^*1118*^ (BDSC #3605) females were used as mating partners. Both males and females were reared at 18 °C and maintained at 25 °C during training. Following training, only the males were shifted to 30 °C for three days to relieve GAL80^ts^-mediated repression prior to dissection. Subsequently, male brains were dissected and stained.

### Training and analysis

During social memory training, the Automated Laser Tracking and Optogenetic Manipulation System (ALTOMS) was utilized to deliver immediate punishment via a 473 nm blue laser when flies violated predefined behavioral contingencies. During the subsequent memory testing phase, the laser remained deactivated, and the inter-fly distance was recorded to calculate the Restraining Index (RI) based on the male’s avoidance behavior.

### Classical conditioning

To systematically optimize the learning effect, four core parameters were modulated: Training Duration (TD), the total duration of a single training session; Bout Length (BL), the fixed duration of each laser activation event; Laser Power (LP), the intensity of the 473 nm stimulus, and Total Punishment duration (TP), the cumulative laser activation throughout the session. In this paradigm, punishment was administered according to a fixed protocol, independent of male behavior. To ensure the overall punishment duration matched that of the operant training condition, the stimulation schedule was derived from the mean TP and bout number quantified during operant sessions using ALTOMS. This generated a regular pattern of alternating laser on/off phases determined as follows:

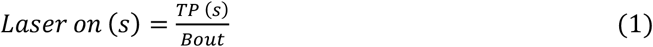

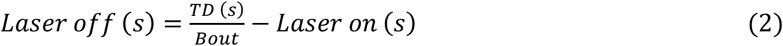

This calculated on/off stimulation cycle was applied continuously from the onset of the session until the end of the training duration (TD). Based on representative operant results (TP = 252 s, bout number = 85) and a TD of 3,600 s, the classical cycle consisted of approximately 3 s laser-on followed by 39 s laser-off phases, repeated consistently throughout the 3,600 s period.

### Operant conditioning

In the operant conditioning paradigm, punishment was contingency-dependent, triggered by the male fly’s proximity to the female. To optimize this learning process, three core parameters were defined: Training Duration (TD), the total duration of the training session; Laser Power (LP), the intensity of the 473 nm stimulus (mW), and Restraining Distance (RD), the spatial threshold (mm) defining the punishment zone. During training, the ALTOMS monitored the real-time distance between the pair. If a male fly remained within the RD of the female for more than 1 s, laser punishment was immediately administered. This stimulus was maintained continuously until the male moved beyond the RD threshold.

### Calculation of behavioral score (Restraining Index, RI)

The Restraining Index (RI) was employed to quantify male avoidance behavior during the testing phase. The RI represents the percentage of time a male remained beyond the predefined proximity threshold from the female over a fixed test duration of 1800 s.

For this study, courtship time was defined as the cumulative duration (in seconds) that the male spent within a 4 mm radius of the female. If copulation occurred, the pair was considered to have remained together for the remainder of the 1800 s test period. The behavioral scores were calculated as follows:

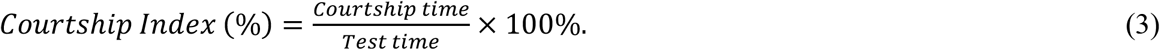

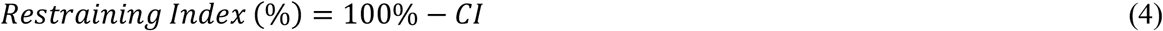

### CSR

The CSR approach utilizes a transfer function to establish a mapping between system outputs and parameter–label (P–L) combinations, thereby characterizing how process parameters affect optimization objectives. The general form of the equation is:

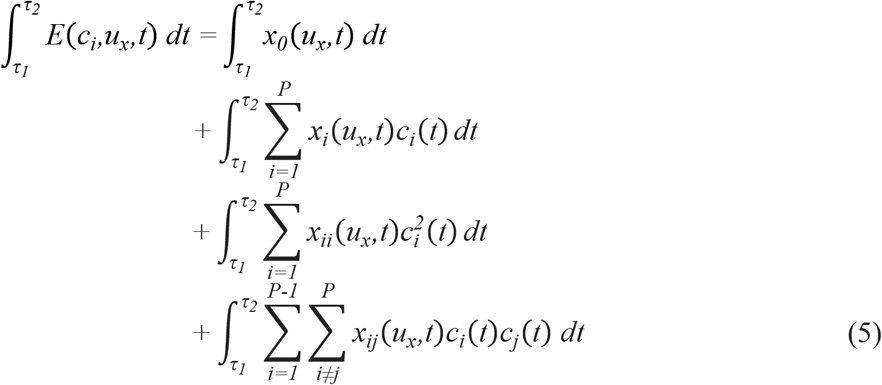

Here, E(c_i_,u_x_,t) denotes the optimization objective (i.e., the system output), c_i_(t) is the i-th process parameter, and P is the total number of parameters considered.

The coefficients x_0_, x_i_, x_ii_, and x_ij_ correspond to the constant, linear, quadratic, and interaction terms, respectively. In total, (P^2^+3P+2)/2 coefficients are required, which can be estimated with far fewer experiments than conventional full-combination testing (P^2^+3P+2)/2≪L^P^. Once the coefficients of the model are fitted, the function can be applied to rapidly identify the optimal P– L combination and its corresponding globally optimized outcome.

To systematically optimize the effect of process parameters on performance, the CSR function was integrated with the Orthogonal-Array-Composite-Design (OACD)^25^. For operant conditioning, the primary parameters optimized were training duration, restraining distance, and laser power. For classical conditioning, optimization focused on training duration, bout length, total punishment duration, and laser power.

For practical implementation, a customized application was developed using MATLAB to support CSR fitting and multi-objective optimization. This tool incorporates three modules: (1) a Least Squares fitting procedure to determine CSR coefficients from P–L combinations and experimental data^26^; (2) prediction of the global optimal P–L configuration using the Interior Point Method applied to the dynamic second-order nonlinear function^27^; and (3) computation of a comprehensive optimization ratio R for performance comparison across conditions. Finally, the coefficient of determination (R^2^) was used to evaluate the goodness of fit and predictive reliability of the CSR model relative to experimental observations.

### RNA sequencing

#### Tissue dissociation

The three adult *Drosophila* brains for each training group were digested with an enzyme cocktail, including dispase (2 U), collagenase I (100 μg/mL), and trypsin-EDTA (0.05%), for 15 minutes at 25 °C with gentle pipetting every 5 minutes. The 10% FBS was then added to the supernatant to stop the enzymatic reaction. The leftover tissue chunks kept digesting with the same enzyme cocktail for 10 min at 25 °C with gentle pipetting every 5 min. Next, the dissociated single cells from two digestions were combined and washed three times with PBS + 0.04% BSA. The viability and cell number were determined and adjusted to a concentration of 1,000 cells/μL for each sample.

#### scRNA-seq library preparation and analysis

The total of 20,000 single cells (1,000 cells/ μL) was collected to construct scRNA-seq libraries using Chromium Next GEM Single Cell 3’ Kit v3.1 (10x Genomics) following the manufacturer’s instructions. The libraries were then sequenced on an Illumina NovaSeq 6000 platform, targeting 20,000 reads per cell. The sequencing reads were mapped to the published date through Cell Ranger v8.0.1 (10x Genomics), and the background and ambient RNA were filtered out using CellBendeR^19^. The subsequent quality control, including (1) unique molecular identifier (UMI) counts, (2) mitochondrial gene content < 20, (3) doublet removal (DoubleFinder v2.0.4), and (4) gene expression level per cell, was done using the Seurat package v.5.1.0 in R. Next, normalization of the processed single-cell datasets was performed using SCTransform and integrated for further gene expression analysis. RunPCA and FindCluster functions in Seurat were used for principal component analysis (PCA) and cell cluster identification. Cell cluster annotation was identified based on published data^19-21^. The FingMarkers function in Seurat was performed to recognize differentially expressed genes (DEGs; p-adjust < 0.05; average log2FC > |0.58|). The clusterProfiler R package (v4.10.1) was utilized to conduct gene ontology (GO) analysis.

### Immunohistochemistry

*Drosophila* brains were first incubated in blocking buffer (10% normal goat serum [NGS], 0.02% NaN_3_, and 2% Triton X-100 in PBS) at 4 °C for 16–18 h. Samples were then transferred to a primary antibody solution (1% NGS, 0.02% NaN_3_, and 0.25% Triton X-100 in PBS) containing either mouse 4F3 anti-DLG (1:20, Hybridoma Bank, University of Iowa) **or** chicken anti-GFP (1:200, Abcam, ab13970), and incubated at 4 °C for 3 days. Following three washes in washing buffer (3% NaCl and 1% Triton X-100 in PBS) for 20 min each, brains were incubated with the corresponding secondary antibody—goat anti-mouse Biotin-XX (1:200, Invitrogen, B-2763) for DLG staining or goat anti-chicken Alexa Fluor 488 (1:200, Abcam, ab150169) for GFP staining—at 4 °C for 2 days. Subsequently, brains were washed three times (20 min each) with wash buffer and, if applicable, incubated with CF633 (1:200, Biotium) at 4 °C for 1 day. Finally, samples were post-fixed in 4% paraformaldehyde (PFA) for 15 min prior to mounting.

### Tissue expansion

Brain tissues were first incubated in a solution containing methacrylic acid N-hydroxysuccinimide ester (MA-NHS) at 4 °C for 16–18 h. Following this step, samples were rinsed twice with PBS at room temperature and then immersed in monomer solution at 4 °C for 5 min. This immersion was repeated twice to allow sufficient diffusion of the monomer prior to gel polymerization.

The tissues were subsequently transferred into a pre-gel solution composed of 2 M sodium chloride (NaCl), 2.5% acrylamide (AA), 0.15% N,N′-methylenebisacrylamide (BIS), 8.625% sodium acrylate (SA), 0.01% 4-hydroxy-TEMPO (4-HT), 0.2% tetramethylethylenediamine (TEMED), and 0.2% ammonium persulfate (APS) in PBS. APS served as the polymerization initiator. Gelation was performed in a custom chamber (∼200 µm thick, ∼9 µL volume) at 37 °C for approximately 2 h.

After gelation, samples were incubated in digestion buffer (50 mM Tris, pH 8.0; 1 mM EDTA; 0.5% Triton X-100; 0.8 M guanidine HCl) supplemented with proteinase K (8 U/mL) and gently agitated at room temperature for 16–18 h. Finally, digested samples were equilibrated in distilled water in a 6-cm dish for at least 1 h, with water exchanged every 20 min, until complete isotropic expansion of the hydrogel-embedded tissues was achieved.

### Spinning disk microscopic imaging

Expanded samples were immersed in double-distilled water for imaging. Image acquisition was performed using a spinning disk confocal microscope (Andor Dragonfly 200, Oxford Instruments, UK) equipped with a 20×/0.5 NA water-immersion objective. Image tiles were subsequently aligned and merged using Imaris Stitcher (v9.9.0, Oxford Instruments, UK). Post-acquisition processing and three-dimensional reconstruction were carried out in Imaris (v9.6.0, Oxford Instruments, UK) to generate high-resolution visualizations suitable for downstream structural and biological analyses.

## Acknowledgements

The authors thank the Brain Research Center, National Tsing-Hua University, Taiwan, for the spinning disk confocal and ALTOMS assistance. This work was also supported by the Brain Research Center under the Higher Education Sprout Project, Taiwan Ministry of Education. We also thank the support from Taiwan National Science and Technology Council.(Grant number: 113-2636-B-007-004 and 113-2311-B-007-013) and National Tsing Hua University.

